# Integrating multi-system environmental factors to predict brain and behavior in adolescents

**DOI:** 10.1101/2024.12.17.628982

**Authors:** Jivesh Ramduny, Samuel Paskewitz, Inti A. Brazil, Arielle Baskin-Sommers

## Abstract

**Objective:** Environmental factors have long been shown to influence brain structure and adolescent psychopathology. However, almost no research has included environmental factors spanning micro-to-macro-systems, brain structure, and psychopathology in an integrated framework. Here, we assessed the ways and degree to which multi-system environmental factors during late childhood predict subcortical volume and psychopathology during early adolescence.

**Method:** We used the baseline and 2-year follow-up data from the Adolescent Brain Cognitive Development^SM^ Study (*N* = 2,766). A Bayesian latent profile analysis was applied to obtain distinct multi-system environmental profiles during late childhood. The profiles were used in a path analysis to predict their direct and indirect effects on subcortical volume and psychopathology during early adolescence.

**Results:** Bayesian latent profile analysis revealed nine environmental profiles. Two distinct profiles predicted greater externalizing problems in adolescents: (i) adversity across, family, school, and neighborhood systems and (ii) family conflict and low school involvement. In contrast, a profile of family and neighborhood affluence predicted fewer externalizing difficulties. Further, family and neighborhood affluence predicted higher subcortical volume, which in turn, predicted fewer externalizing problems; whereas, family economic and neighborhood adversity predicted lower subcortical volume, which in turn, predicted greater externalizing difficulties.

**Conclusion:** We captured direct and indirect influences of environmental factors across multiple systems on externalizing psychopathology. Specifying the equifinal pathways to externalizing psychopathology serves to provide an evidence base for establishing different types of interventions based on the needs and risk profiles of youth.

**Diversity and Inclusion Statement:** The current study is part of the ongoing Adolescent Brain Cognitive Development^SM^ Study (ABCD Study®) for which youth are recruited from elementary schools in the United States that are informed by gender, race, ethnicity, socioeconomic status, and urbanicity. The ABCD Study® aims to recruit youth longitudinally by sampling the sociodemographic makeup of the US population. Two of the authors self-identifies as a member of one or more historically underrepresented racial and/or ethnic groups in science. One of the authors identifies as a part of an underrepresented gender group in science. The authors also are representative of the communities for which data was collected and contributed to design, analysis, and/or interpretation of the work. Finally, every effort was made to cite the work of authors from underrepresented and minoritized groups in academic research.

## Introduction

An extensive body of research has identified that adolescent psychopathology relates to changes in brain development and experiences in the environment. However, much of this work has been siloed into work specifying the environmental or neurobiological factors related to psychopathology adolescence.

On the one hand, decades of research have documented that environmental systems influence the development of psychopathology (i.e., externalizing and internalizing problems)^1–5^. Meta-analyses report medium-to-large effects between adversity in adolescents’ families (e.g., conflict, caregiver nonacceptance) and neighborhoods (e.g., experiencing violence or disadvantage) and adolescent psychopathology^6,7^. Externalizing problems have been associated with adverse experiences in the form of family poverty, harsh parenting, association with deviant peers, concentrated disadvantage, and exposure to community violence^2^. Similarly, internalizing problems have been linked to maternal depression, maltreatment by a caregiver, peer victimization, and exposure to community violence can exacerbate internalizing difficulties in youth^3–5^. On the other hand, neurobiological theories of adolescent psychopathology have emphasized that structural changes, a general indicator of brain health. In particular, subcortical brain volumes, are sensitive to externalizing and internalizing problems during adolescent development^8,9^. These brain regions support self-regulation and affective processing^10^, and differences in their structural volumes have been related to various aspects of adolescent psychopathology^11^.

Integrating environmental and neurobiological factors, some researchers have shown that experiences within different environmental systems influence structural brain development^12^. Adverse experiences—such as low household income, harsh parenting, maltreatment, peer victimization, neighborhood disadvantage, and community violence—have shown to be associated with reduced subcortical volumes in regions that are involved in cognitive and emotion processing^13–16^. For example, children from lower income families and those exposed to community violence have smaller hippocampal and amygdala volumes^13–16^.

Additionally, maternal harsh parenting has been related to smaller amygdala volume^17^ and childhood maltreatment has been associated with smaller hippocampal volume^18^. Researchers also have shown that anti-poverty policies in the United States mitigate socioeconomic-related brain differences, such that in high cost of living states with generous anti-poverty programs, the association between family income and hippocampal volume resembled that of lower cost of living states^14^. However, the majority of these studies have focused on examining the influences of single environmental systems on brain structures, and have not examined the combined influence of family, school, neighborhood, and policy factors.

Moreover, many of these studies have been designed with relatively small sample sizes, more restricted sociodemographic samples (e.g., all low income, all high income, all maltreatment, all disadvantaged neighborhood), and a focus on selective subcortical regions (e.g., hippocampus and amygdala).

Research on the influence of environmental experiences on adolescent psychopathology, brain on psychopathology, and their interactions (albeit in limited ways) has laid a strong foundation for understanding how psychopathology may unfold in different contexts for different youth. However, there is a need to employ an integrated approach^19,20^ that fully captures the transactions among multiple environmental systems, brain development, and psychopathology. First, we must do better in estimating the multiple, and often interacting, environmental systems youth encounter—some that relate to a youth’s immediate surroundings (e.g., family, school, neighborhood)^21^ while others that relate to larger societal and cultural contexts (e.g., policies, law)^22^. Second, we need to understand how interconnected environmental experiences, *directly* or *indirectly*, relate to subcortical brain structure and psychopathology.

In the present study, we tested the relationships among multiple environmental systems, subcortical gray matter (GM) volume, and psychopathology using the Adolescent Brain Cognitive Development^SM^ Study (ABCD Study®)^23^. First, we aimed to identify distinct profiles of youth during late childhood from <u>family</u>, <u>school</u>, <u>neighborhood</u>, and <u>policy</u> systems using a novel Bayesian latent profile analysis (LPA)^24^ method. Most often, researchers use multiple regression and factor analysis with single, or possibly two, environmental systems and subcortical ROIs, limiting their ability to capture complex (within-person) interactions of multi-system environmental factors and their associations with brain structures. LPA is an analytic approach that derive unique profiles of individuals that exhibit similar characteristics (e.g., environmental experiences). Here, we used the Bayesian LPA method as it has been shown to capture more nuanced and more certain profiles than conventional LPA^24^. We hypothesized the emergence of latent profiles characterized by moderate and high adversities across multiple environmental systems. Given the novelty of combining multi-system environments in the Bayesian LPA method, we did not have specific hypotheses about latent profiles that could capture more subtle variations in adversities.

Second, we aimed to assess the ways and degree to which the multi-system environmental profiles during late childhood predict subcortical GM volume and psychopathology (i.e., externalizing, internalizing) during early adolescence. Based on prior research^1–5,13–16^, we hypothesized that profiles describing high adverse family and neighborhood environments would predict smaller subcortical GM volume and greater externalizing problems. Following evidence that brain structures can mediate the association between poverty and externalizing problems during adolescence^16,25^, we also expected an indirect relationship between adverse multi-system environments and externalizing via subcortical GM volume.

## Methods

### Participants

The ABCD Study is a ten-year longitudinal study that tracks the development of children and adolescents across 21 research sites in the United States^23^. We used the demographic, environmental, behavioral, and imaging data from the ABCD Study Release 5.0 (*N*=11,868) at baseline (9-10 years) and 2-year follow-up (11-12 years) (https://abcdstudy.org/). The ABCD Study obtained approval from a centralized Institutional Review Board (IRB) located at the University of California, San Diego in addition to obtaining local IRB approval from each of the imaging sites. Written assent was provided by the youth and written informed consent was obtained by their parents or guardians. Only participants who had complete demographic, environmental, psychopathology, and imaging data in addition to passing successfully the MRI data quality control criteria described by the ABCD Data Analysis and Informatics Center (DAIC)^26^ (https://wiki.abcdstudy.org/release-notes/imaging/quality-control.html) were included in the study. A total of 2,766 youth remained in the study (**Table 1**).

**Table 1.**
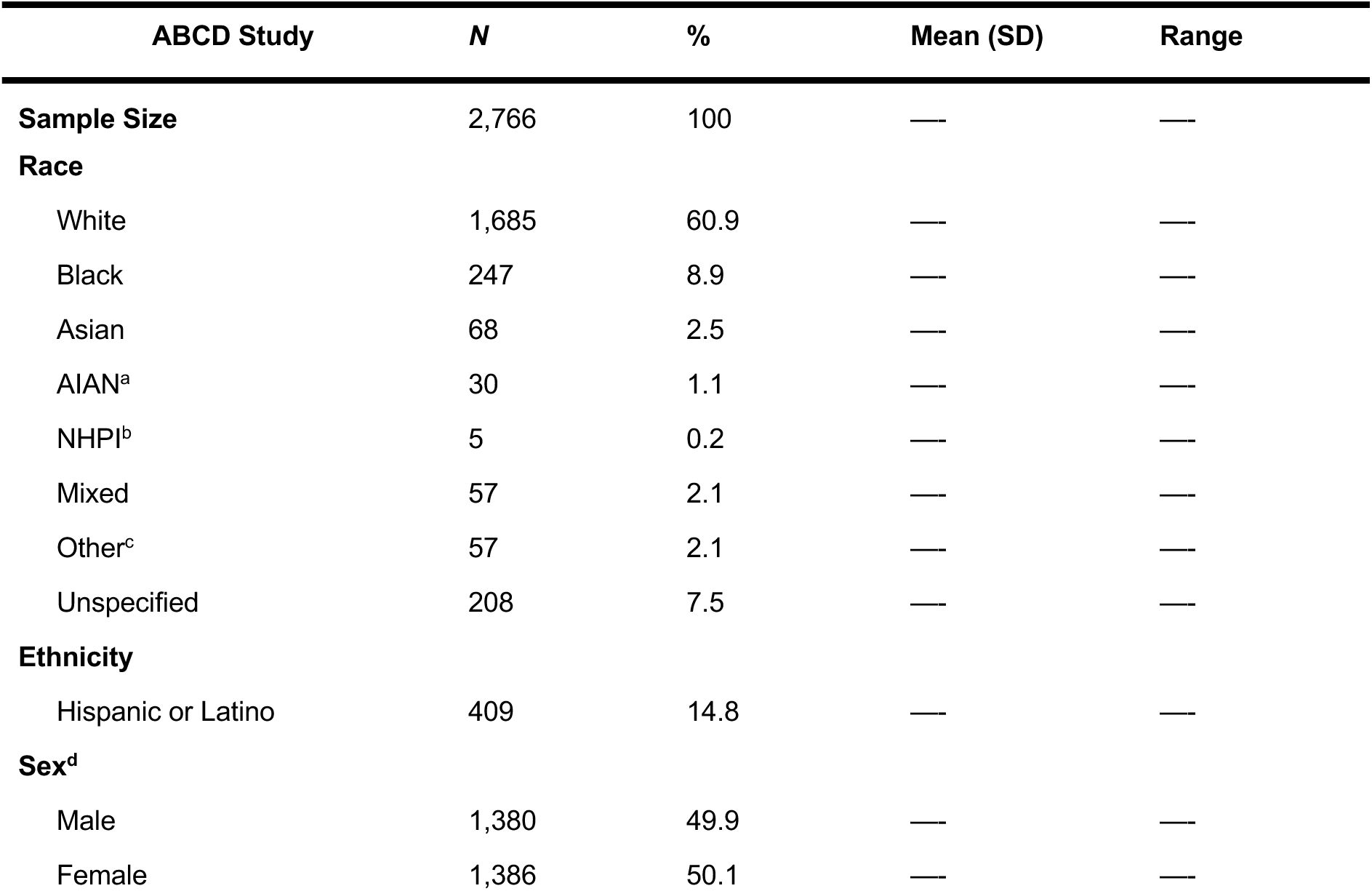
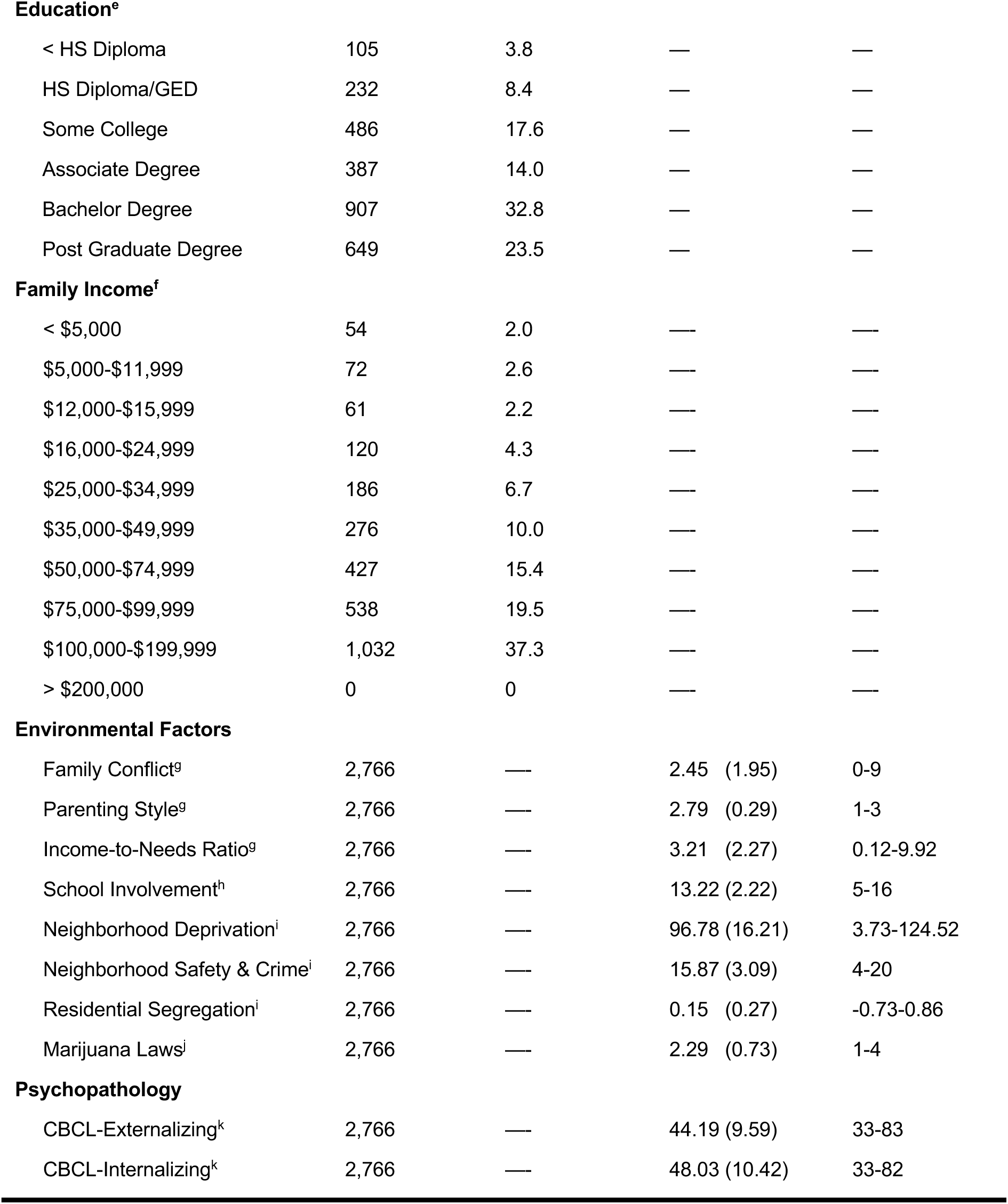
Demographic, environmental, and behavioral characteristics derived from the ABCD Study NIMH Data Archive Release 5.0. ^a^AIAN = American Indian and Alaska Native. ^b^NHPI = Native Hawaiian and Pacific Islander. ^c^Other race/ethnicity corresponds to Eastern and Western European, Afro-Carribean/Indo-Carribbean/West Indian, Middle Eastern/North African in addition to parents who selected “Other race” to indicate that the predefined groups did not apply to them. ^d^Participant sex denotes youth’s sex assigned at birth. ^e^Education refers to the highest grade or level of school a parent has completed or the highest degree they have received. ^f^Family Income refers to the total income in a household and the income bands are provided by the ABCD Study. ^g^Family-related environmental factors describe family conflict, parenting style, and income-to-needs ratio at baseline. ^h^School-related environmental factor describes school involvement at baseline. ^i^Neighborhood-related environmental factors describe neighborhood deprivation, neighborhood crime and safety, and residential segregation at baseline. ^j^Policy-related environmental factor refers to marijuana laws with regards to cannabis legalization in the United States at baseline. ^k^CBCL = Child Behavior Checklist indexing externalizing and internalizing psychopathology at 2-year follow-up. Note that the CBCL externalizing and internalizing scores were ***T***-standardized. We also compared the characteristics of our sample with the full ABCD sample to test for differences in demographics, multi-system environments, and psychopathology (**Supplement 1**).

| ABCD Study | *N* | % | Mean (SD) | Range |
| --- | --- | --- | --- | --- |
| **Sample Size** | 2,766 | 100 | — | — |
| **Race** |  |  |  |  |
| White | 1,685 | 60.9 | — | — |
| Black | 247 | 8.9 | — | — |
| Asian | 68 | 2.5 | — | — |
| AIAN<sup>a</sup> | 30 | 1.1 | — | — |
| NHPI<sup>b</sup> | 5 | 0.2 | — | — |
| Mixed | 57 | 2.1 | — | — |
| Other<sup>c</sup> | 57 | 2.1 | — | — |
| Unspecified | 208 | 7.5 | — | — |
| **Ethnicity** |  |  |  |  |
| Hispanic or Latino | 409 | 14.8 | — | — |
| **Sex<sup>d</sup>** |  |  |  |  |
| Male | 1,380 | 49.9 | — | — |
| Female | 1,386 | 50.1 | — | — |
| **Education<sup>e</sup>** |  |  |  |  |
| < HS Diploma | 105 | 3.8 | — | — |
| HS Diploma/GED | 232 | 8.4 | — | — |
| Some College | 486 | 17.6 | — | — |
| Associate Degree | 387 | 14.0 | — | — |
| Bachelor Degree | 907 | 32.8 | — | — |
| Post Graduate Degree | 649 | 23.5 | — | — |
| **Family Income<sup>f</sup>** |  |  |  |  |
| < \$5,000 | 54 | 2.0 | — | — |
| \$5,000-\$11,999 | 72 | 2.6 | — | — |
| \$12,000-\$15,999 | 61 | 2.2 | — | — |
| \$16,000-\$24,999 | 120 | 4.3 | — | — |
| \$25,000-\$34,999 | 186 | 6.7 | — | — |
| \$35,000-\$49,999 | 276 | 10.0 | — | — |
| \$50,000-\$74,999 | 427 | 15.4 | — | — |
| \$75,000-\$99,999 | 538 | 19.5 | — | — |
| \$100,000-\$199,999 | 1,032 | 37.3 | — | — |
| > \$200,000 | 0 | 0 | — | — |
| **Environmental Factors** |  |  |  |  |
| Family Conflict<sup>g</sup> | 2,766 | — | 2.45 (1.95) | 0-9 |
| Parenting Style<sup>g</sup> | 2,766 | — | 2.79 (0.29) | 1-3 |
| Income-to-Needs Ratio<sup>g</sup> | 2,766 | — | 3.21 (2.27) | 0.12-9.92 |
| School Involvement<sup>h</sup> | 2,766 | — | 13.22 (2.22) | 5-16 |
| Neighborhood Deprivation<sup>i</sup> | 2,766 | — | 96.78 (16.21) | 3.73-124.52 |
| Neighborhood Safety & Crime<sup>i</sup> | 2,766 | — | 15.87 (3.09) | 4-20 |
| Residential Segregation<sup>i</sup> | 2,766 | — | 0.15 (0.27) | -0.73-0.86 |
| Marijuana Laws<sup>j</sup> | 2,766 | — | 2.29 (0.73) | 1-4 |

**Table 1. Demographic, environmental, and behavioral characteristics derived from the ABCD Study NIMH Data Archive Release 5.0.** <sup>a</sup>AIAN = American Indian and Alaska Native. <sup>b</sup>NHPI =
|  |  |  |  |  |
| --- | --- | --- | --- | --- |
| CBCL-Externalizing<sup>k</sup> | 2,766 | — | 44.19 (9.59) | 33-83 |
| CBCL-Internalizing<sup>k</sup> | 2,766 | — | 48.03 (10.42) | 33-82 |

### Environmental Data

The environmental factors were obtained at baseline from the ABCD Study Culture and Environment^27^, Linked External Data^28^, and Adolescent Neural Urbanome^29^ batteries. We focused on multi-system environmental factors that have shown relationships with subcortical GM volume and youth psychopathology^1–5,13–16^. Choosing environmental factors across multiple systems from the ABCD Study is challenging as they show some degree of correlation with each other. Further, the Bayesian LPA method that we used to identify distinct profiles of youth requires that the factors follow a normal distribution^24^. Therefore, environmental factors that had binary outcomes were not included in this study. Policy data available in the ABCD Study, such as medicaid expansion, naloxone policies, and Good Samaritan law were binary outcomes and had no variability in their distributions^28^. By contrast, the marijuana laws variable was not binary, and it displayed sufficient variability within the ABCD sample.

#### Family Conflict

The Family Conflict subscale from the Family Environment Scale (FES) consists of nine items, indicating whether each statement is True or False by the youth for most family members. The nine items were summed and reverse scored to obtain a measure of family conflict for each youth. A higher value on the FES Conflict subscale indicates less family conflict perceived by the youth.

#### Parenting Style

The Acceptance Scale from the Children’s Reports of Parental Behavior Inventory (CRPBI) consists of five items, where the youth describe their caregivers’ parenting style on a 3-point scale. The mean score from the five items was used as a measure of parenting style for each youth. A higher value on the CRPBI Acceptance Scale indicates warmer parenting style perceived by the youth.

#### Income-to-needs Ratio

The income-to-needs ratio was calculated as the median of the income band described by the ABCD Study divided by the federal poverty level based on the respective household size.

The median of the first income band was set at $5,000 and the median for the last income band was set at $200,000. The federal poverty level was obtained from the Department of Health and Human Services (https://www.healthcare.gov/glossary/federal-poverty-level-fpl/).

An income-to-needs ratio of 1 indicates living at the poverty threshold, a ratio greater than 1 denotes living above the poverty threshold, and a ratio less than 1 denotes living below the poverty threshold.

#### School Involvement

The School Involvement subscale from the School Risk and Protective Factors (SRPF) questionnaire contains four items as indicators of positive involvement in school. The scores from the four items were summed to obtain a measure of school involvement for each youth. A higher value on the SRPF School Involvement subscale indicates more school involvement perceived by the youth.

#### Neighborhood Deprivation

The area deprivation index (ADI) reflects the weighted sum of 17 composite scores related to employment, education, income and poverty, and housing using the youth’s home address from the 2011-2015 American Community Survey (**Table S1**, available online). We reverse coded the ADI scores with higher ADI values indicating lower neighborhood disadvantage.

#### Neighborhood Safety and Crime

The Safety from Crime item scales from the PhenX Toolkit describe three statements administered to the parent related to feeling safe walking in their neighborhood, violence in their neighborhood, and crime in their neighborhood. Each item is rated on a 5-point Likert scale ranging from “strongly agree (5)” to “strongly disagree (1)”. Only one statement was administered to youth to assess their feelings about safety and crime in their neighborhood. The scores from the parent and youth item scales were summed to obtain a measure of neighborhood safety and crime for each youth. A higher value on the Safety from Crime item scales indicates a safer neighborhood perceived by the parent and youth.

#### Residential Segregation

The index of concentration at the extremes (ICE) was used to examine the extent to which a population in a specified area is concentrated into the wealthiest and poorest extremes of a specified social distributions. ICE measures the distributions of affluence and poverty within racial or ethnic groups across the wealthiest and poorest areas of the community based on the 2014-2018 American Community Survey. It ranges from -1 to 1 such that a positive value indicates concentration of a racial/ethnic group in affluent areas, a negative value denotes concentration of a racial/ethnic group in impoverished areas, and a value closer to 0 indicates no concentration of a racial/ethnic group in either affluent or poorer areas of the community.

#### Marijuana Laws

Currently, there are 38 states that have legalized cannabis for medical use, 24 states that provide legal access to cannabis for recreational use, and 9 states that allow low THC, high CBD products either for medical purposes or as a legal defense in the United States (http://www.ncsl.org/research/health/state-medical-marijuana-laws.aspx). States that legalize either recreational or medical cannabis use reflect more liberal marijuana laws as opposed to those which forbid legal access to cannabis, therefore being more conservative. We obtained policy data representing marijuana laws across the United States, and reverse coded the categories for cannabis legalization. Cannabis legalization was categorized as follows: 1:No legal access to cannabis; 2:Low THC/CBD; 3:Medical; and 4:Recreational. A higher value in the cannabis legalization categories indicates a less conservative law.

### Psychopathology Data

The Child Behavior Checklist (CBCL) is a parent-report assessment that was used to measure externalizing and internalizing behaviors^30^. We used the parent-report as youth self-report on externalizing and internalizing were not available for the 2-year follow-up. Externalizing behaviors tend to reflect symptoms such as aggression and rule-breaking, whereas internalizing symptoms capture anxiety and depression. The CBCL externalizing and internalizing scores were obtained from their respective syndrome scales that were subsequently ***T***-standardized. The higher the CBCL externalizing and internalizing ***T***-scores, the greater the risk of experiencing behavioral and emotional problems.

### MRI Data Acquisition

Structural T1-weighted and T2-weighted MRI scans were acquired using Siemens Prisma, Philips, and GE 750 3T scanners with a 32-channel head coil^23^. 3D MPRAGE T1-weighted and 3D FSE T2-weighted volumes with spatial resolution 1×1×1mm^3^ were obtained for each youth at 2-year follow-up. The structural MRI data were preprocessed using the DAIC standard processing pipeline^26^.

### Subcortical Gray Matter Volume

For each participant, the subcortical GM structures were labeled using an automated, atlas-based, volumetric segmentation procedure from FreeSurfer^26^. These structures include 19 ROIs corresponding to bilateral nucleus accumbens, amygdala, caudate, hippocampus, pallidum, putamen, thalamus, ventral diencephalon, cerebellum, and midline brainstem.

### Bayesian Latent Profile Analysis

We performed a Bayesian latent profile analysis (LPA) using the environmental factors. LPA explains a set of indicator variables by grouping participants into latent profiles, i.e., categories of individuals with similar characteristics. The LPA model assumes that each participant belongs to a single latent profile and that indicator variables have independent normal likelihoods with means that vary across profiles^24^. Conventional methods for determining the correct number of latent profiles often give conflicting results for which number of profiles is best to select. We used a Bayesian form of LPA based on the Dirichlet process mixture model^62^ that automatically detects the correct number of latent profiles. This Bayesian LPA method was implemented using the Python package *vbayesfa*^24^. We used proportional reduction in classification entropy to assess the distinctiveness of the profiles inferred.

### Integrated Approach

We examined relationships among multi-system environmental profiles, subcortical GM volume, and externalizing and internalizing using a path analysis (**Figure 1**). The path analysis was conducted by including only one sibling, i.e., one child per family, to limit family dependency confounds, with the *lavaan* package in R version 4.3.2. We tested the direct and indirect effects of the multi-system environmental profiles simultaneously—a <u>direct</u> effect represents a relationship between a multi-system environmental profile and subcortical GM volume *or* between a multi-system environmental profile and externalizing/internalizing whereas an <u>indirect</u> effect represents a relationship between a multi-system environmental profile and externalizing/internalizing *via* the subcortical GM volume. For direct effects, the coefficient estimates, standardized errors, and statistical significance were reported. For indirect effects, the coefficient estimates and 95% confidence intervals (CIs), which were obtained from a bootstrapping procedure, were reported.

**Figure 1.**
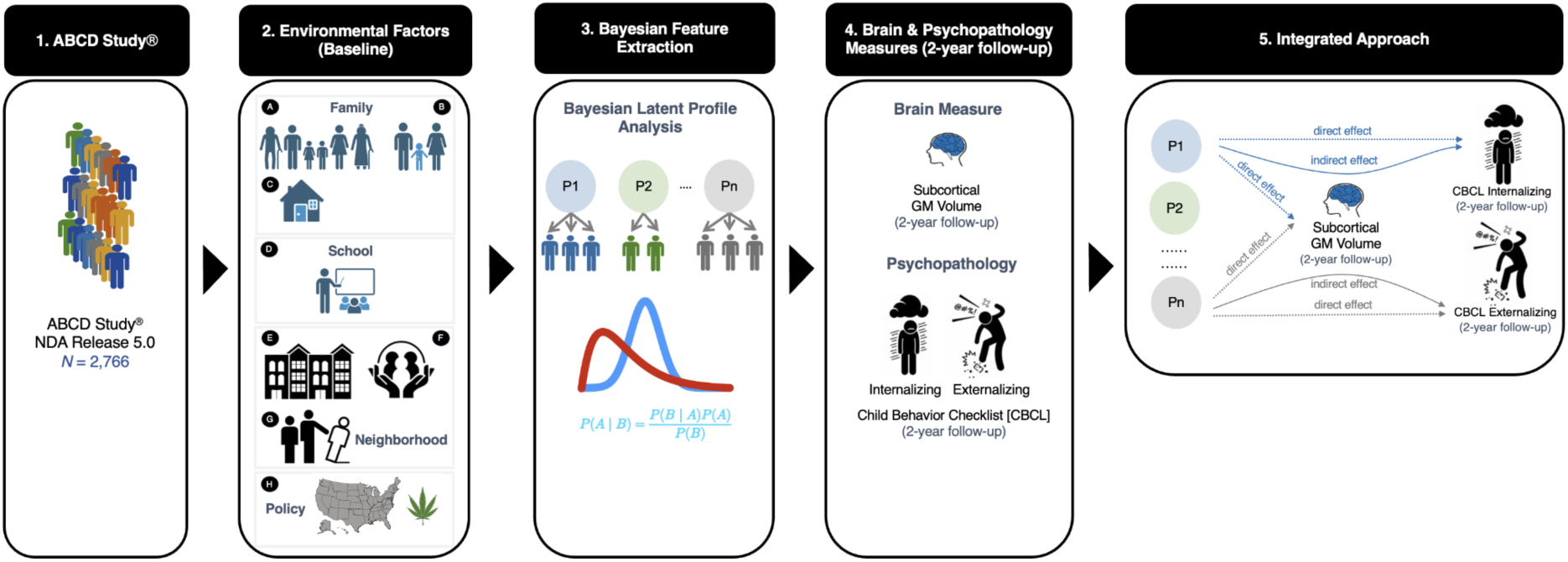
Integrated Approach. 1. The ABCD Study® NDA Release 5.0 was used to obtain environmental, brain, and behavioral data from late childhood (baseline) to early adolescence (2-year follow-up). 2. The environmental factors were obtained at baseline and they captured multiple systems including family, school, neighborhood, and policy. Family-related factors correspond to (A) family conflict, (B) parenting style, and (C) income-to-needs ratio. School-related factor corresponds to (D) school involvement. Neighborhood-related factors correspond to (E) neighborhood disadvantage, (F) neighborhood safety and crime, and (G) residential segregation. Policy-related factor corresponds to (H) marijuana laws with regards to cannabis legalization in the United States. 3. The environmental factors were then used to generate distinct profiles (denoted by P1, P2, …., Pn) of youth automatically that share similar characteristics without *a priori* specifying the number of latent profiles using a Bayesian latent profile analysis framework. We assigned each participant to their most probable profile and then treated the profiles like an observed variable. 4. Subsequently, the subcortical gray matter (GM) volume was obtained at 2-year follow-up from the ABCD Study. The subcortical GM volume corresponds to 19 ROIs including the bilateral nucleus accumbens, amygdala, caudate, hippocampus, pallidum, putamen, thalamus, ventral diencephalon, cerebellum, and extending to the midline brainstem. The behavioral measures also were obtained from the Child Behavior Checklist (CBCL) which correspond to externalizing and internalizing psychopathology at 2-year follow-up. 5. The integrated approach represents a path analysis linking the multi-system environmental factors (denoted by P1, P2, …., Pn), subcortical GM volume, and externalizing and internalizing psychopathology. A <u>direct</u> effect captures the relationship between a multi-system environmental profile and subcortical GM volume *or* between a multi-system environmental profile and externalizing/internalizing behavior. An <u>indirect</u> direct effect captures the relationship between a multi-system environmental profile and externalizing/internalizing behavior *via* the subcortical GM volume.

## Results

### Description of the multi-system environmental profiles

The Bayesian LPA model produced 9 distinct profiles with an excellent proportional reduction in entropy of 0.90 (**Figure 2**). In creating labels, we wanted to use judgment-free language and avoid listing factors as positive or negative. Further, given the array of factors representing environmental systems, it was difficult to come up with a labeling scheme that would apply appropriately to all factors. Therefore, we opted to use descriptive labels for each profile based on the collection of factors that deviated from the mean. For each profile, the characteristics of all environmental factors are provided in **Table 2**.

**Figure 2.**
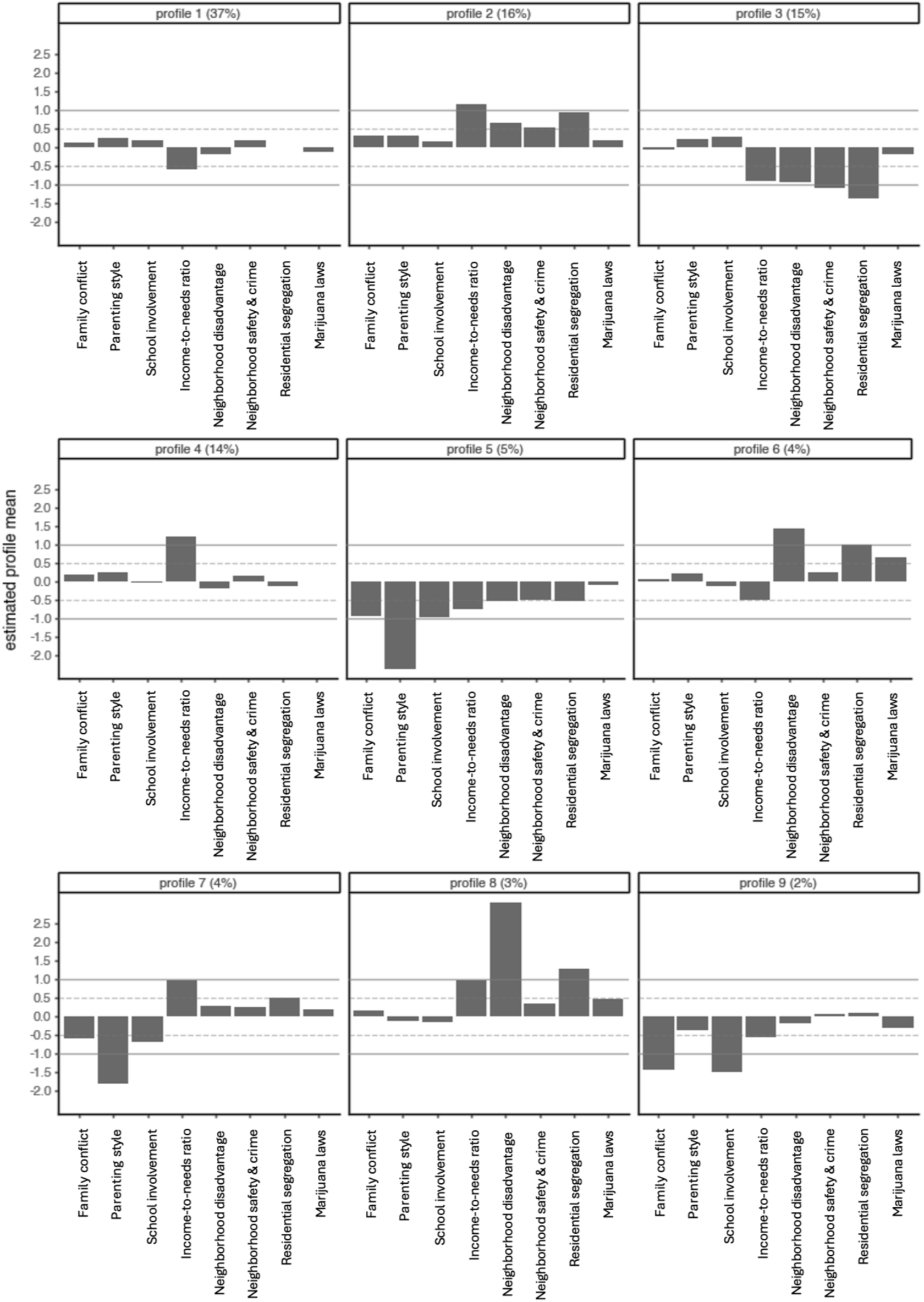
Descriptions of the multi-system environmental profiles obtained from family, school, neighborhood, and policy factors. Family factors correspond to family conflict, parenting style, and income-to-needs ratio. The school factor corresponds to school involvement. Neighborhood factors correspond to neighborhood disadvantage, neighborhood safety and crime, and residential segregation. The policy factor corresponds to marijuana laws. The proportion of youth in each distinct profile also is displayed. Note that family conflict, neighborhood disadvantage, and marijuana laws have been reverse coded. Dotted gray line denotes responses 0.50±SD from the mean of each profile. Solid gray line denotes responses 1.0±SD from the mean of each profile.

**Table 2.** Characteristics of the multi-system environmental profiles. For each profile, the mean household size and mean income band defined by the ABCD Study are shown. The ABCD income range reflects the mean income band for a given profile. Based on 2024 U.S. Federal Poverty Guidelines issued by the Department of Health and Human Services (https://aspe.hhs.gov/topics/poverty-economic-mobility/poverty-guidelines), income ranges for Profiles 3 and 5 fall below the federal poverty line for a household of 5 people. The federal poverty line for a household of 5 people is $36,580 in 48 contiguous states and the District of Columbia. For the remaining profiles, the income ranges do not fall below the federal poverty line for their respective mean household sizes. For each profile, the mean and range of the individual environmental factors also are shown.

| Profile | Mean Household Size | Mean Income Band | ABCD Income Range | Family Conflict (FES) | Parenting Style (CRPBI) | School Involvement (SRPF) | Neighborhood Disadvantage (ADI) | Neighborhood Safety&Crime (PhenX) | Residential Segregation (ICE) | Marijuana Laws (mj) |
| --- | --- | --- | --- | --- | --- | --- | --- | --- | --- | --- |
| 1 | 5 | 6 | \$35,000-\$49,000 | 1.74 (0-9) | 2.86 (2.2-3.0) | 13.6 (6-16) | 99.9 (72.2-116) | 16.4 (6-20) | 0.155 (-0.39-0.63) | 2.38 (1-4) |
| 2 | 5 | 9 | \$100,000-\$199,999 | 1.38 (0-8) | 2.89 (2.4-3.0) | 13.6 (6-16) | 85.5 (60.0-102) | 17.6 (7-20) | 0.424 (0.11-0.74) | 2.13 (1-3) |
| 3 | 5 | 5 | **\$25,000-\$34,999** | 2.10 (0-9) | 2.86 (2.2-3.0) | 13.9 (7-16) | 113 (87.4-125) | 12.2 (4-20) | -0.246 (-0.73-0.09) | 2.44 (1-4) |
| 4 | 4 | 9 | \$100,000-\$199,999 | 1.69 (0-8) | 2.86 (2.4-3.0) | 13.1 (6-16) | 101 (69.8-119) | 16.3 (5-20) | 0.095 (-0.44-0.41) | 2.29 (1-4) |
| 5 | 5 | 5 | **\$25,000-\$34,999** | 4.00 (0-9) | 2.05 (1.0-2.4) | 11.0 (4-16) | 106 (68.0-125) | 14.1 (5-20) | -0.000004 (-0.54-0.53) | 2.37 (1-4) |
| 6 | 5 | 6 | \$35,000-\$49,999 | 1.88 (0-8) | 2.86 (2.2-3.0) | 12.8 (8-16) | 70.3 (40.8-90.0) | 16.6 (7-20) | 0.441 (-0.21-0.74) | 1.74 (1-3) |
| 7 | 5 | 8 | \$75,000-\$99,999 | 3.12 (0-9) | 2.21 (1.0-2.6) | 11.4 (4-16) | 91.4 (55.3-115) | 16.6 (11-20) | 0.309 (-0.13-0.69) | 2.15 (1-3) |
| 8 | 4 | 8 | \$75,000-\$99,999 | 1.59 (0-8) | 2.77 (1.2-3.0) | 12.9 (8-16) | 40.8 (3.73-61.7) | 17.0 (12-20) | 0.534 (0.08-0.86) | 1.91 (1-3) |
| 9 | 6 | 6 | \$35,000-\$49,999 | 5.56 (2-9) | 2.69 (2.4-3.0) | 8.79 (5-13) | 99.8 (83.6-116) | 16.3 (6-20) | 0.183 (-0.31-0.54) | 2.57 (1-3) |

**Figure 3.**
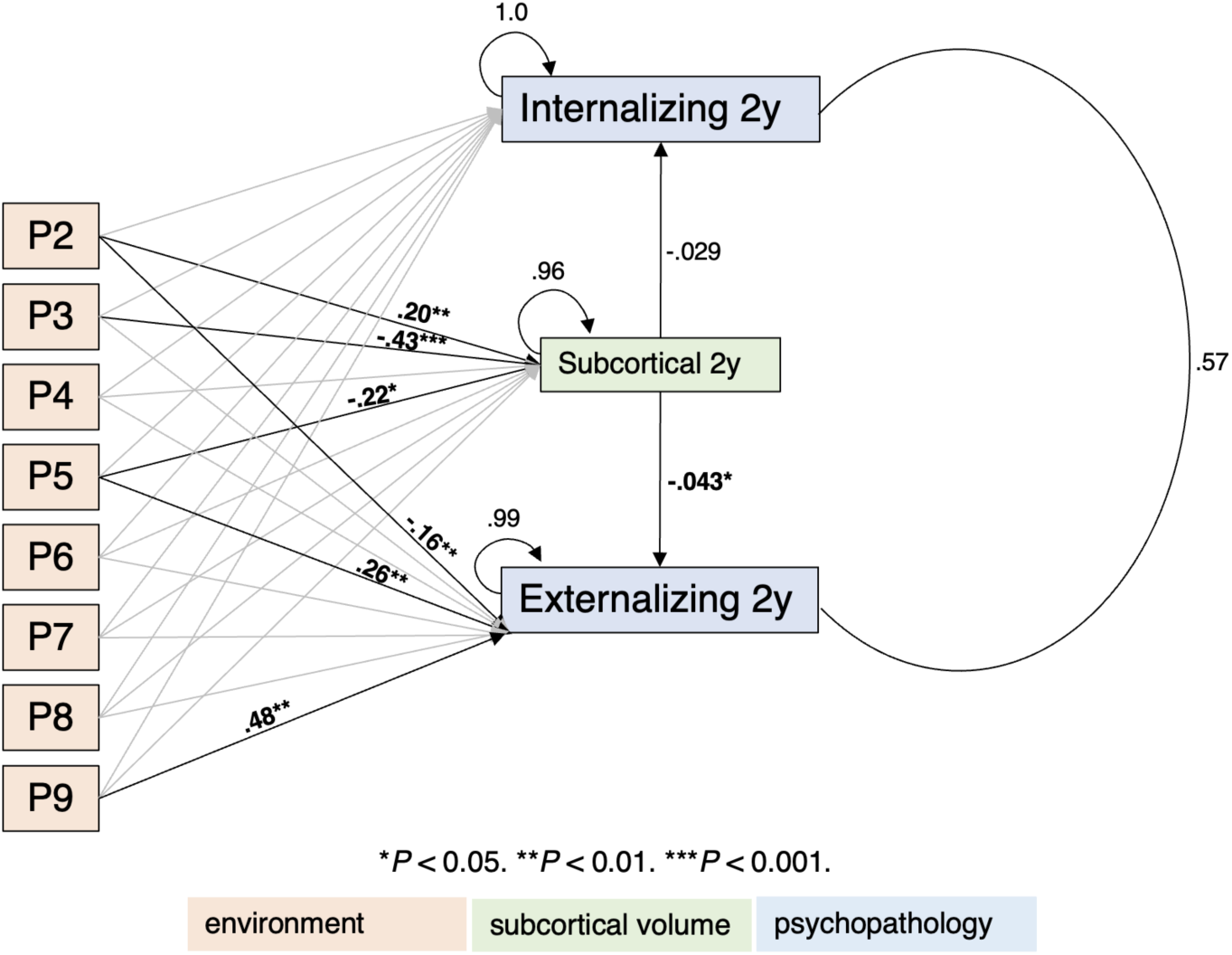
Integrated approach linking multi-system environmental profiles, subcortical GM volume, and externalizing/internalizing psychopathology. The integrated approach was operationalized using a path analysis from structural equation modeling. The multi-system environmental profiles (denoted by P2, …., P9) are derived from family, school, neighborhood, and policy factors at baseline. The imaging measure corresponds to the subcortical gray matter (GM) volume at 2-year follow-up. Externalizing and internalizing psychopathology are derived from the Child Behavior Checklist (CBCL) at 2-year follow-up. Bold black arrows represent the significant relationships (direct and indirect) with their respective coefficient estimates. Gray arrows indicate the relationships between the multi-system environmental profiles and subcortical GM volume or externalizing/internalizing psychopathology that are not significant. Circle arrows capture the respective variances of subcortical GM volume, externalizing psychopathology, and internalizing psychopathology. Circular line denotes the correlation between externalizing and internalizing psychopathology. A bootstrapping procedure was performed to estimate the 95% confidence intervals (CIs) of the indirect effects by selecting samples of youth randomly to perform the path analysis over 10,000 times. \**P* < 0.05. \*\**P* < 0.01. \*\*\**P* < 0.001.

**Profile 1** was characterized by below average family income [mean income band=6] representing 37% of the sample. **Profile 2** was characterized by family [mean income band=9] and neighborhood affluence 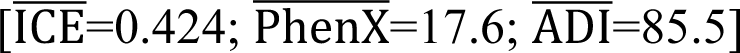 representing 16% of the sample. **Profile 3** was characterized by family economic [mean income band=5] and neighborhood adversity 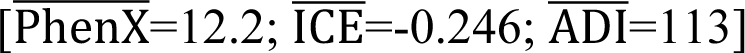 representing 15% of the sample. **Profile 4** was characterized by family economic affluence [mean income band=9] representing 14% of the sample. **Profile 5** was characterized by adversity across family 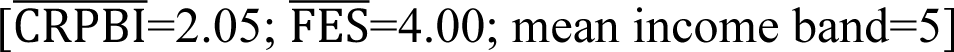, school 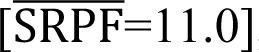, and neighborhood systems 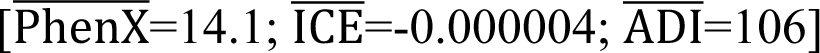 representing 5% of the sample. **Profile 6** was characterized by neighborhood affluence 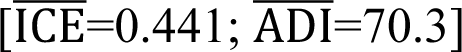 and liberal marijuana policy 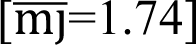, but below average family income [mean income band=6] representing 4% of the sample. **Profile 7** was characterized by adverse family interactions 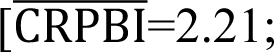 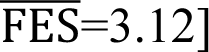 and low school involvement 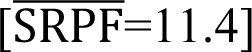, but family economic [mean income band=8], and neighborhood affluence 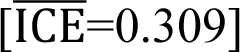 representing 4% of the sample. **Profile 8** was characterized by neighborhood 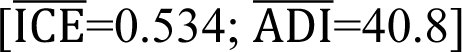 and family economic affluence [mean income band = 8], with somewhat liberal marijuana policy 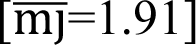 representing 3% of the sample. **Profile 9** was characterized by family conflict 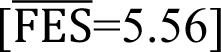 and low school involvement 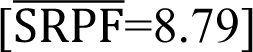 representing 2% of the sample. The multi-system environmental profiles differed significantly in the distributions of participant race/ethnicity, US region, and sex assigned at birth (**Figure S1**, available online).

### Integrated Approach

The nine multi-system environmental profiles were dummy coded and Profile 1 was treated as the reference profile as youth recorded average responses for the majority of the environmental factors. First, we observed <u>direct</u> effects of the multi-system environmental profiles on subcortical GM volume. **Profile 2**, i.e., family and neighborhood affluence during late childhood, compared to Profile 1 predicted higher subcortical GM volume during early adolescence (estimate [SE]=0.20 [0.059], *P*=0.001). **Profile 3**, i.e., family economic and neighborhood adversity during late childhood, compared to Profile 1 predicted lower subcortical GM volume during early adolescence (estimate [SE]=-0.43 [0.059], *P*<0.001).

**Profile 5**, i.e., adversity across, family, school, and neighborhood systems during late childhood, compared to Profile 1 predicted lower subcortical GM volume during early adolescence (estimate [SE]=-0.22 [0.094], *P*=0.018).

Next, we observed <u>direct</u> effects of the multi-system environmental profiles on externalizing psychopathology. **Profile 2**, compared to Profile 1 predicted fewer externalizing problems during early adolescence (estimate [SE]=-0.16 [0.060], *P*=0.009). **Profile 5**, compared to Profile 1, predicted greater externalizing difficulties during early adolescence (estimate [SE]=0.26 [0.095], *P*=0.006). **Profile 9**, i.e., family conflict and low school involvement during late childhood, compared to Profile 1 predicted greater externalizing problems during early adolescence (estimate [SE]=0.48 [0.14], *P*=0.001).

We also observed <u>indirect</u> effects of the multi-system environmental factors on externalizing psychopathology *via* subcortical GM volume. The path **Profile 2→Subcortical 2y→Externalizing 2y** indicated that family and neighborhood affluence during late childhood predicted higher subcortical GM volume during early adolescence, which in turn, predicted fewer externalizing problems (estimate=-0.006, 95% bootstrap CI=[-0.020, - 0.00032]). The path **Profile 3→Subcortical 2y→Externalizing 2y** indicated that family economic and neighborhood adversity during late childhood predicted lower subcortical GM volume, which in turn, predicted greater externalizing difficulties during early childhood (estimate=0.012, 95% bootstrap CI=[0.0012, 0.036]). See supplemental material for tests of robustness by treating baseline subcortical GM volume and participant sex assigned at birth, respectively (**Figures S2-S3**, available online).

## Discussion

The goal of the present study was to capture complex interactions among multiple environmental systems and examine the impact of these systems on youth’s brain and behavioral development as they transition from childhood to adolescence. To achieve this goal, we first applied a novel Bayesian LPA framework to identify distinct subgroups of children from family, school, neighborhood, and policy systems. We then assessed the ways and degree to which the multi-system environmental profiles predicted subcortical GM volume and externalizing and internalizing during early adolescence. The Bayesian LPA revealed nine distinct profiles with excellent certainty and discrimination. The integrated approach further captured direct and indirect influences of the multi-system environmental profiles on subcortical GM volume and externalizing. Together, the transactions among multi-system environments, brain structure, and psychopathology revealed an equifinality in the ways subcortical volume and externalizing difficulties may be influenced during adolescence.

Many foundational theories of development emphasize the importance of examining multiple contexts at different levels of proximity to youth^21,22,31,32^. Some researchers take the approach of documenting the additive accumulation of multiple environmental risks as it relates to adolescent psychopathology^33,34^. Increasingly, though, researchers are identifying the co-occurring interplay among different environmental experiences to identify relative contributions of these environments on adolescent psychopathology^35,36^. A recent study using the ABCD Study provided evidence of four distinct profiles of perceived threat across family, school, and neighborhood systems^35^. These profiles reflected low threat across the three systems, elevated threat in the neighborhood, elevated threat in the family, and elevated threat across the three systems. Youth in the elevated threat across all systems profile had poorer mental health outcomes, but youth in the family threat profile uniquely showed more disruptive behavior symptoms and youth in the elevated neighborhood threat profile displayed increased sleep problems. Another study found distinct groups of youth who experienced low, medium, and high adversity, and maternal depression from family and neighborhood systems^36^. Youth reported lowest externalizing and internalizing symptoms in the low adversity followed by medium, maternal depression, and high adversity profiles.

These profile analyses are a great step in identifying the interplay among environmental systems, and their impact on psychopathology. However, traditional LPA methods impose a trade-off between number of participants in a sample and number of profiles that produces meaningful discrimination—a large number of profiles leads to harder interpretation whereas a smaller number of profiles leads to relative levels of low, medium, and high responses. Our Bayesian LPA approach overcomes traditional limitations, revealing reliable profiles with subtle variations in childhood environmental experiences^24^ that align with U.S. census data^37^ and highlight regional disparities in poverty and income (e.g., **Profile 3** reflects the South, with the highest childhood poverty and 1 in 5 Black women in poverty, while **Profile 2** reflects the West, where White families dominate higher income levels). This approach advances our understanding of how diverse environmental experiences interact and offers a more nuanced framework for identifying risk and protective factors in adolescent development.

We found three pathways from which late childhood multi-system environmental profiles *directly* influenced subcortical volume. First, youth experiencing family and neighborhood affluence predicted larger subcortical volume compared to youth in a below average family income profile. Generally, environments that provide more opportunities and quality resources can act as a buffer against stress for children and relate to better physical health compared to those who belong to lower socioeconomic families and disadvantaged neighborhoods^38^. Second, youth experiencing family economic and neighborhood adversity had smaller subcortical volume compared to youth living in an environment with below average family income. Third, adversity across, family, school, and neighborhood systems during late childhood predicted smaller subcortical volume compared to lower family income alone. The latter two effects are consistent with animal research showing that certain subcortical regions are susceptible to effects of early adversities^39^. Early life adversity negatively impacts neurogenesis (process of generating new neurons) in the hippocampus and has been associated with increased dendritic arborization (process by which neurons create connections with other neurons) in the amygdala^39^. In human research, children living in or near poverty exhibit structural brain differences^15,16^, including smaller subcortical volumes.

Combined with previous research, the present study documents the relative effect that access to fewer economic resources (at the family and neighborhood level) has on brain health. This highlights the need to implement policies related to anti-poverty programs and neighborhood resources (e.g., safety, amenities) to facilitate healthy developmental trajectories for youth.

The present analysis also revealed the unique ways different environmental systems *directly* influence adolescent externalizing psychopathology. Youth who experienced relative family and neighborhood affluence compared to youth living in families with lower income showed fewer externalizing symptoms. However, youth who experienced adversity across family, school, and neighborhood systems or who experienced family conflict and low school involvement during late childhood had more externalizing problems in adolescence. It is well-documented that youth who experience multi-system adversity show more externalizing problems^40,41^. For instance, the family stress model posits that poverty, unsafe neighborhoods, and economic instability stress parents, undermining their emotional resources, and leading to greater family conflict, and eventually harsher parenting (and child externalizing problems). This model has been supported by a host of empirical work^42–44^, highlighting that broader environments play a role in shaping the proximal environment for the developing youth. Notably, though, experiences of family conflict and low school involvement reflected another robust risk pathway for externalizing, suggesting for some youth adversity localized to family and school interactions confer risk for externalizing.

Parenting that is harsh and inconsistent has been linked robustly to the development of externalizing behaviors^45,46^. This type of parenting is thought to model aggression for young people, to undermine their ability to develop emotion regulation skills necessary for related constructs such as empathy, and inconsistency in parenting is thought to lead to reward contingencies that make aggression or breaking rules useful in some contexts. Further, youth who experience conflict at home often bring this history to school where they begin these types of cycles with peers and teachers, leading to trouble in school and often social rejection. These findings underscore the need for researchers and clinicians to assess risk across multiple systems. The conceptualization of a youth’s behavior should be based on the combinations of risk factors that are influential for *that* person, providing a more personalized and targeted approach to intervention.

Critically, the integrated path analysis highlighted key pathways whereby experiences within environmental systems predicted externalizing problems via subcortical GM volume. Family and neighborhood affluence during late childhood predicted higher subcortical GM volume during early adolescence, which in turn, predicted fewer externalizing problems. This pathway reveals how relative affluence buffers against externalizing psychopathology via GM volume in brain regions responsible for reward, sensorimotor, cognitive, and emotion processing. In contrast, family economic and neighborhood adversity predicted lower subcortical GM volume, which in turn, predicted greater externalizing difficulties. This finding is consistent with a previous study showing that subcortical GM trajectories mediate the relationships between preschool socioeconomic status and high-risk behaviors^15^.

Therefore, there is growing evidence to show that socioeconomic adversities can negatively impact structural brain development, which may increase the risk of mental health problems through stress and lack of experiences that enrich development^16,47^. However, subcortical GM volume did not mediate the relationship between the profile that described the most adverse experiences across multiple systems (**Profile 5**) and externalizing problems. When a child experiences multiple strong adversities, these adverse experiences provide a “push” to a given outcome such that the importance of biological factors in these environments might be diminished^48^. This is supported by the social push hypothesis^48^—when there are more and stronger environmental pushes (e.g., harsh parenting in the context of neighborhood disadvantage and crime) towards externalizing problems (such as aggression), biological risk for aggression may matter less. Our findings support the consideration of an integrated approach to parse the *direct* and *indirect* transactions between multiple environmental systems, subcortical GM volume, and externalizing psychopathology.

Before concluding, it is important to note some limitations. First, we focused on subcortical GM volume as subcortical structures are sensitive to socioeconomic resources and psychopathology during childhood and adolescence^1–5,13–16^. However, other brain measures also have been associated with socioeconomic resources^49,50^. Future work could identify pathways that facilitate transactions between multiple environments, multi-modal brain measures, and psychopathology. Second, while we used the ABCD Study as it samples the sociodemographic variations longitudinally in the US population, our sample may not capture the full spectrum of adversities that youth experience. As there was no relationship between the multi-system environmental profiles and internalizing problems, future work could extend the environmental factors that may confer risk for internalizing psychopathology. Finally, this study follows a time lagged design to assess the ways and degree to which multi-system environmental factors during late childhood *predict* subcortical volume and psychopathology during early adolescence. We cannot infer the ways and extent to which subcortical volume and psychopathology *change* longitudinally from childhood to adolescence. Future work assessing how environmental systems influence baseline subcortical volume and externalizing across different developmental stages could fully test the transactional nature of environment-brain-psychopathology.

In conclusion, subcortical brain development and adolescent psychopathology are influenced by multiple environmental systems related to family, school, neighborhood, and policy factors. Our integrated approach highlights multiple equifinal pathways to adolescent externalizing psychopathology, some directly via the environment and others via structural brain development. Adverse environmental experiences should not just be viewed as “challenges”, but instead as experiences that can shape the brain and behavior, ultimately impacting mental health. Measuring environmental and neural factors could give us more information about the status of particular risk/protective factors and help us refine our understanding of *who* benefits from *what* interventions.

## Acknowledgments

Data used in the preparation of this article were obtained from the ABCD Study® (<u>abcdstudy.org/</u>), held in the NIMH Data Archive (NDA). This is a multisite, longitudinal study designed to recruit more than 10,000 children aged 9-10 and follow them over 10 years into early adulthood. The ABCD Study is supported by the National Institutes of Health (NIH) and additional federal partners under award numbers: U01DA041048, U01DA050989, U01DA051016, U01DA041022, U01DA051018, U01DA051037, U01DA050987, U01DA041174, U01DA041106, U01DA041117, U01DA041028, U01DA041134, U01DA050988, U01DA051039, U01DA041156, U01DA041025, U01DA041120, U01DA051038, U01DA041148, U01DA041093, U01DA041089, U24DA041123, U24DA041147.

The full list of federal supporters is available at https://abcdstudy.org/federal-partners.html. The complete lists of participating sites and study investigators can be found at https://abcdstudy.org/consortium_members/. The ABCD Consortium investigators designed and implemented the study and/or provided the data but did not necessarily participate in the analysis or writing of this report. This manuscript reflects the views of the authors and may not reflect the opinions or views of the NIH or ABCD Consortium investigators. Additional support for this work was made possible from NIEHS R01-ES032295, R01-ES031074, and R21DA057592. This work also obtained support from the Yale Kavli Institute for Neuroscience and the Wu Tsai Institute at Yale University. We thank the Yale Center for Research Computing for guidance and use of the research computing infrastructure.

## Data Availability

The ABCD Study® is openly available following access permission granted to one or multiple NIMH Data Archive (NDA) Collections (https://nda.nih.gov/nda/access-data-info). The ABCD data repository grows and changes over time (https://nda.nih.gov/). The ABCD data used in this report came from the tabulated data which can be navigated from the Data Dictionary Explorer Release 5.1 (https://data-dict.abcdstudy.org/<u>?</u>). The detailed descriptions of the assessments used in this report are available from the ABCD Study Protocols (https://abcdstudy.org/scientists/protocols/).

## Code Availability

The analysis code for the Bayesian LPA framework can be found at https://github.com/SamPaskewitz/dpm.lpa. The analysis code for the integrated approach can be found at https://github.com/JRam02/embeddedbrain.

## Author Contributions Statement

J.R curated the demographic, environmental, behavioral, and imaging data from the ABCD Study®. S.P developed the Bayesian LPA framework. J.R and S.P developed and tested the integrated approach. J.R, S.P, and A.B.-S conceptualized this study, interpreted the analyses, and wrote the original draft of the manuscript. J.R, S.P, I.A.B, and A.B.-S reviewed and edited the manuscript. A.B.-S supervised this study.

