## Supplementary Material for "Integrating multi-system environmental factors to predict brain and behavior in adolescents"

RH: Predicting brain and behavior via multi-system factors

Jivesh Ramduny, Ph.D., Samuel Paskewitz, Ph.D., Inti A. Brazil, Ph.D., Arielle Baskin-Sommers, Ph.D.

Drs. Ramduny, Paskewitz, and Baskin-Sommers are with the Department of Psychology, Yale University. Dr. Ramduny also is with the Kavli Institute for Neuroscience, Yale University. Dr. Baskin-Sommers also is with the Department of Psychiatry and Wu Tsai Institute, Yale University. Dr. Brazil is with the Donders Institute for Brain, Cognition and Behaviour, Radboud University. Dr. Brazil also is with the Forensic Psychiatric Centre Pompestichting, The Netherlands.

Samuel Paskewitz, Ph.D., served as the statistical expert for this research.

**Acknowledgments**
Data used in the preparation of this article were obtained from the ABCD Study® (abcdstudy.org/), held in the NIMH Data Archive (NDA). This is a multisite, longitudinal study designed to recruit more than 10,000 children aged 9-10 and follow them over 10 years into early adulthood. The ABCD Study is supported by the National Institutes of Health (NIH) and additional federal partners under award numbers: U01DA041048, U01DA050989, U01DA051016, U01DA041022, U01DA051018, U01DA051037, U01DA050987, U01DA041174, U01DA041106, U01DA041117, U01DA041028, U01DA041134, U01DA050988, U01DA051039, U01DA041156, U01DA041025, U01DA041120, U01DA051038, U01DA041148, U01DA041093, U01DA041089, U24DA041123, U24DA041147. The full list of federal supporters is available at https://abcdstudy.org/federal-partners.html. The complete lists of participating sites and study investigators can be found at https://abcdstudy.org/consortium_members/. The ABCD Consortium investigators designed and implemented the study and/or provided the data but did not necessarily participate in the analysis or writing of this report. This manuscript reflects the views of the authors and may not reflect the opinions or views of the NIH or ABCD Consortium investigators. Additional support for this work was made possible from NIEHS R01-ES032295, R01-ES031074, and R21DA057592. This work also obtained support from the Yale Kavli Institute for Neuroscience and the Wu Tsai Institute at Yale University. We thank the Yale Center for Research Computing for guidance and use of the research computing infrastructure.

**Disclosure**
Drs. Ramduny, Paskewitz, Brazil, and Baskin-Sommers have reported no biomedical financial interests or potential conflicts of interest.

**Correspondence**

| Variable No. | Variable Name | Variable Description |
|---|---|---|
| 1 | reshist_addr1_adi_edu_l | Percentage of population aged >= 25 years with < 9 years of education |
| 2 | reshist_addr1_adi_edu_h | Percentage of population aged >= 25 years with at least a high school diploma |
| 3 | reshist_addr1_adi_work_c | Percentage of employed persons aged >= 16 years in white collar occupations |
| 4 | reshist_addr1_adi_income | Median family income |
| 5 | reshist_addr1_adi_in_dis | Income disparity defined by Singh (2003) as the log of 100 x ratio of the number of households with < 10000 annual income to the number of households with > 50000 annual income |
| 6 | reshist_addr1_adi_home_v | Median home value |
| 7 | reshist_addr1_adi_rent | Median gross rent |
| 8 | reshist_addr1_adi_mortg | Median monthly mortgage |
| 9 | reshist_addr1_adi_home_o | Percentage of owner |
| 10 | reshist_addr1_adi_crowd | Percentage of occupied housing units with >1 person per room (crowding) |
| 11 | reshist_addr1_adi_unemp | Percentage of civilian labor force population aged >= 16 years unemployed (unemployment rate) |
| 12 | reshist_addr1_adi_pov | Percentage of families below the poverty level |
| 13 | reshist_addr1_adi_b138 | Percentage of population below 138% of the poverty threshold |
| 14 | reshist_addr1_adi_sp | Percentage of single |
| 15 | reshist_addr1_adi_ncar | Percentage of occupied housing units without a motor vehicle |
| 16 | reshist_addr1_adi_ntel | Percentage of occupied housing units without a telephone |
| 17 | reshist_addr1_adi_nplumb | Percentage of occupied housing units without complete plumbing (log) |

**Table S1. Area Deprivation Index.** Variable name and description of census-tract estimates for the area deprivation index (ADI) included in the ABCD Study. The variable names and descriptions are derived from Fan and colleagues.

**Supplement 1**

**Comparing our sample (*N* = 2,766) with the larger ABCD sample (*N* = 5,994)**

Bonferroni correction was applied to correct for multiple comparisons across the demographic, environmental, and psychopathology variables. The *P* values were adjusted based on the number of statistical tests carried out to compare the characteristics of our sample with those from the larger ABCD sample.

**Demographics**

No difference

Participant race/ethnicity: Chi-square test, $\chi^2 = 14.88$, $P = 0.85$

Participant sex assigned at birth: Chi-square test, $\chi^2 = 4.93$, $P = 0.36$

Parental education: Chi-square test, $\chi^2 = 9.39$, $P = 1.0$

Imaging site: Chi-square test, $\chi^2 = 0$, $P = 1.0$

**Environments**

Difference

Marijuana laws: Chi-square test, $\chi^2 = 16.30$, $P = 0.014$

Note: The residuals indicated that the observed count was higher than expected for three categories (i.e., recreational, medical, no legal access) in the larger ABCD sample. The observed count was higher than expected for one category only (i.e., low THC/CBD) in our sample.

No difference

Family conflict: Mann-Whitney test, U = 8423343.50, $P = 1.0$

Parenting style: Mann-Whitney test, U = 8211872.50, $P = 1.0$

Income-to-needs ratio: Mann-Whitney test, U = 8369175.00, $P = 1.0$

School involvement: Mann-Whitney test, U = 8075703.00, $P = 0.69$

Neighborhood disadvantage: Mann-Whitney test, U = 8454448.50, $P = 1.0$

Neighborhood safety & crime: Mann-Whitney test, U = 8228359.50, $P = 1.0$

Residential segregation: Mann-Whitney test, U = 7999363.50, $P = 0.11$

**Psychopathology**

No Difference

CBCL-Externalizing: Mann-Whitney test, U = 8567152.50, $P = 0.15$

CBCL-Internalizing: Mann-Whitney test, U = 8333150.50, $P = 1.0$

**Characteristics of the multi-system environmental profiles**

We examined the characteristics of the multi-system environmental profiles based on participant race/ethnicity, US region, and sex assigned at birth (**Figure S1**, available online). First, there was a significant difference in the distribution of participant race/ethnicity across the 9 profiles (Chi-square test, $\mathcal{X}^2 = 636.70$, $P < 0.001$). While White youth dominated 8 out of 9 profiles, Black youth were more frequent in Profile 3, which was characterized by family economic and neighborhood adversity. Second, we categorized the 21 imaging sites of the youth into four geographical locations based on the census region of the ABCD Study site: Northeast, Midwest, South, and West. There was a significant difference in the distribution of US regions across the 9 profiles (Chi-square test, $\mathcal{X}^2 = 209.04$, $P < 0.001$). Youth who were recruited from Western locations dominated 7 profiles (Profiles 1, 2, 4, 6, 7, 8, 9) followed by youth who belonged to Southern (2 profiles; Profiles 3, 5) geographical areas. Last, there was a significant difference in the distribution of participant sex assigned at birth across the 9 profiles (Chi-square test, $\mathcal{X}^2 = 19.05$, $P = 0.015$). While male youth were more representative in 6 profiles (Profiles 2, 4, 5, 6, 7, 9), female youth were more frequent in the remaining 3 profiles (Profiles 1, 3, 8).

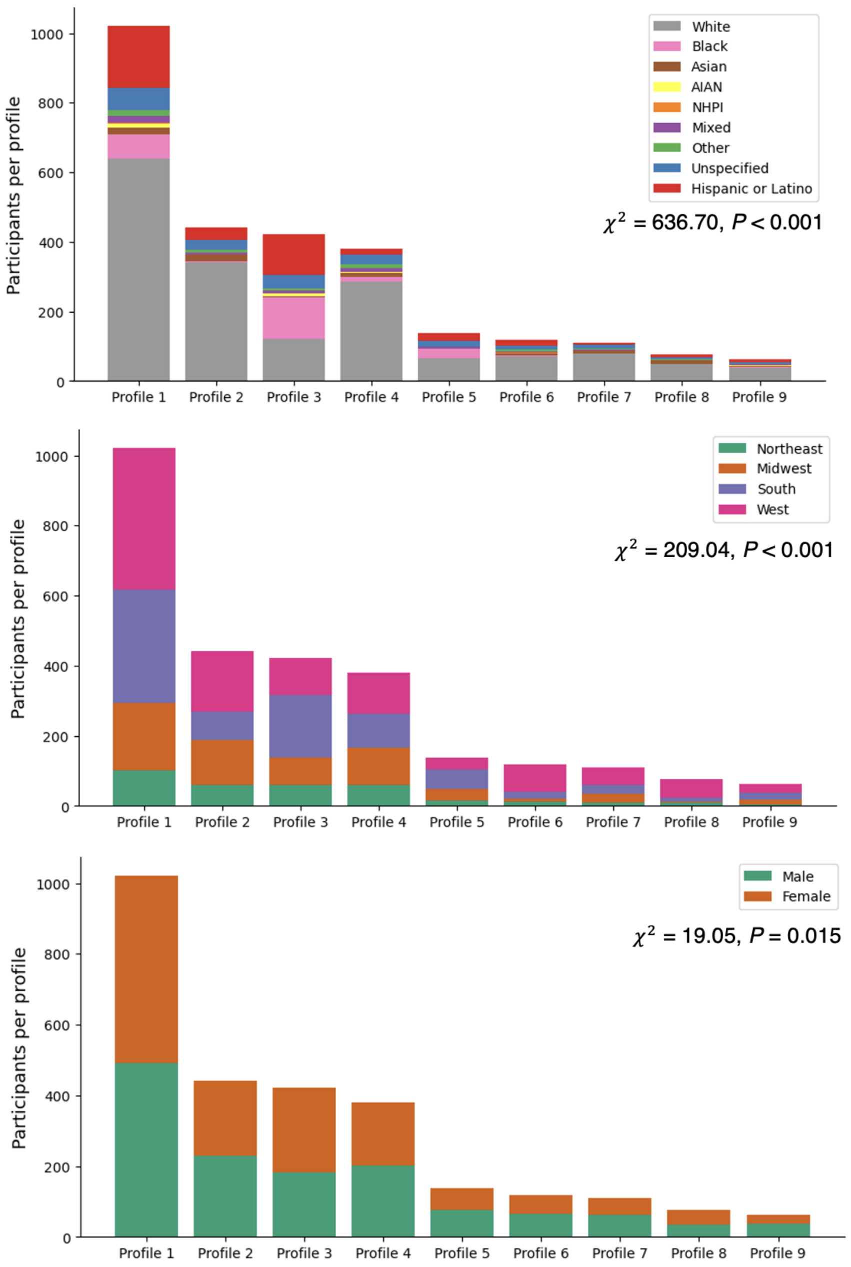


**Figure S1. Characteristics of the multi-system environmental profiles based on participant race/ethnicity, US region, and sex assigned at birth.** Top Panel. Participant race/ethnicity represents White, Black, Hispanic, Asian, American Indian and Alaska Native (AIAN), Native Hawaiian and Pacific Islander (NHPI), Mixed, Other, Unspecified, Hispanic or Latino youth. Youth belonging to Other races/ethnicities correspond to European, African, and multiple races. Middle Panel. The 21 imaging sites are categorized into four geographical locations based on the census region of the ABCD Study® site. *Northeast*: University of Rochester, University of Vermont, University of Pittsburgh, Yale University; *Midwest*: University of Minnesota, University of Michigan, University of Wisconsin-Milwaukee, Washington University in St. Louis; S*outh*: Florida International University, Laureate Institute for Brain Research, Medical University of South Carolina, University of Florida, Virginia Commonwealth University, University of Maryland at Baltimore; and *West*: Children's Hospital Los Angeles, University of Colorado Boulder, SRI International, UCLA, UC San Diego, University of Utah; Oregon Health & Science University. Bottom Panel. Participant sex assigned at birth represents male and female youth.

**Robustness of the integrated approach**

We tested the robustness of the integrated approach by treating baseline subcortical GM volume and participant sex assigned at birth as covariates, respectively (**Figures S2-S3**, available online). When baseline subcortical GM volume was included as an additional covariate, several direct paths between multi-system environmental profiles and subcortical GM volume (i.e., Profile 3→Subcortical 2y) and between multi-system environmental profiles and externalizing (i.e., Profile 2→Externalizing 2y; Profile 5→Externalizing 2y; Profile 9→Externalizing 2y) remained. An indirect effect of the multi-system environmental factors on externalizing via subcortical GM volume (i.e., Profile 3→Subcortical 2y→Externalizing 2y) also remained present. When participant sex assigned at birth was treated as a covariate, all the direct paths between multi-system environmental profiles and subcortical GM volume (i.e., Profile 2→Subcortical 2y; Profile 3→Subcortical 2y; Profile 5→Subcortical 2y) and between multi-system environmental profiles and externalizing (i.e., Profile 2→Externalizing 2y; Profile 5→Externalizing 2y; Profile 9→Externalizing 2y) remained. Similarly, the two indirect effects of the multi-system environmental factors on externalizing via subcortical GM volume (i.e., Profile 2→Subcortical 2y→Externalizing 2y; Profile 3→Subcortical 2y→Externalizing 2y) remained significant.

***Adding baseline subcortical GM volume as a covariate in the integrated approach***

Direct effects between multi-system environmental profiles and subcortical GM volume
**P3→Subcortical 2y: estimate [SE] = -0.029 [0.011], *P* = 0.007**
P4→Subcortical 2y: estimate [SE] = -0.025 [0.011], *P* = 0.025

Direct effects between multi-system environmental profiles and externalizing psychopathology
**P2→Externalizing 2y: estimate [SE] = -0.16 [0.060], *P* = 0.009**
**P5→Externalizing 2y: estimate [SE] = 0.26 [0.095], *P* = 0.006**
**P9→Externalizing 2y: estimate [SE] = 0.48 [0.14], *P* = 0.001**

Indirect effects between multi-system environmental profiles and externalizing psychopathology via subcortical GM volume
**P3→Subcortical 2y→Externalizing 2y: estimate = 0.0012, 95% bootstrap CI = [0.000012, 0.0030]**

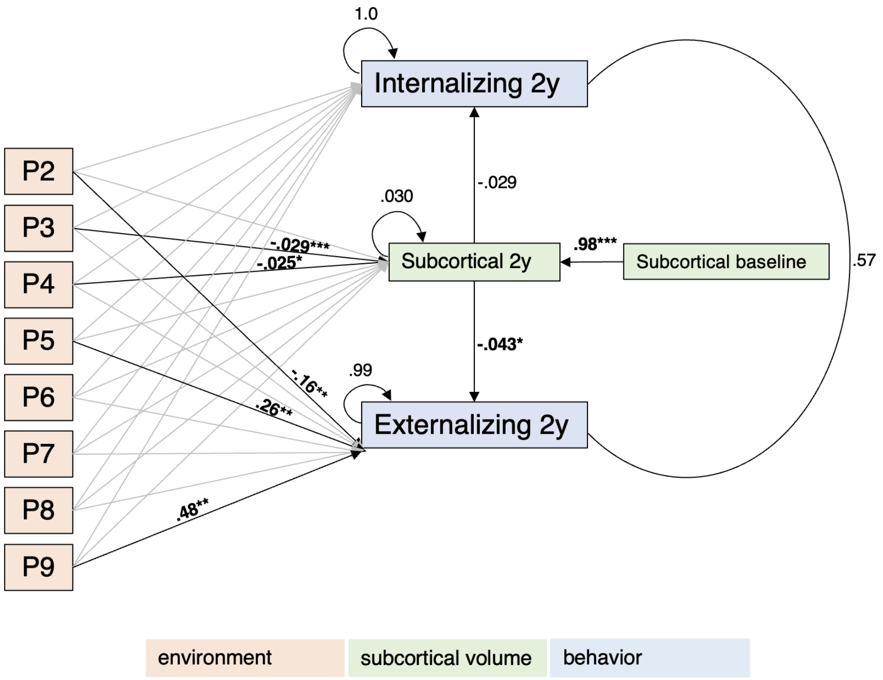


**Figure S2. Integrated approach linking multi-system environmental profiles, subcortical GM volume, and externalizing/internalizing psychopathology.** The integrated approach was operationalized using a path analysis from structural equation modeling and baseline subcortical GM volume was treated as a covariate. The multi-system environmental profiles (denoted by P2, …., P9) are derived from family, school, neighborhood, and policy factors at baseline. The imaging measure corresponds to the subcortical gray matter (GM) volume at 2-year follow-up. Externalizing and internalizing psychopathology are derived from the Child Behavior Checklist (CBCL) at 2-year follow-up. Bold black arrows represent the significant relationships (direct and indirect) with their respective coefficient estimates. Gray arrows indicate the relationships between the multi-system environmental profiles and subcortical GM volume or externalizing/internalizing psychopathology that are not significant. Circle arrows capture the respective variances of subcortical GM volume, externalizing psychopathology, and internalizing psychopathology. Circular line denotes the correlation between externalizing and internalizing psychopathology. A bootstrapping procedure was performed to estimate the 95% confidence intervals (CIs) of the indirect effects by selecting samples of youth randomly to perform the path analysis over 10,000 times. *$P < 0.05$. **$P < 0.01$. ***$P < 0.001$.

***Adding participant sex assigned at birth as a covariate in the integrated approach***

<u>Direct effects between multi-system environmental profiles and subcortical GM volume</u>
**P2→Subcortical 2y: estimate [SE] = 0.17 [0.053], *P* = 0.001**
**P3→Subcortical 2y: estimate [SE] = -0.40 [0.053], *P* < 0.001**
**P5→Subcortical 2y: estimate [SE] = -0.29 [0.084], *P* = 0.001**

<u>Direct effects between multi-system environmental profiles and externalizing psychopathology</u>
**P2→Externalizing 2y: estimate [SE] = -0.16 [0.060], *P* = 0.010**
**P5→Externalizing 2y: estimate [SE] = 0.24 [0.095], *P* = 0.012**
**P9→Externalizing 2y: estimate [SE] = 0.46 [0.14], *P* = 0.001**

<u>Indirect effects between multi-system environmental profiles and externalizing psychopathology via subcortical GM volume</u>

**P2→Subcortical 2y→Externalizing 2y: estimate = -0.087, 95% bootstrap CI = [-0.027, -0.004]**
**P3→Subcortical 2y→Externalizing 2y: estimate = 0.033, 95% bootstrap CI = [0.015, 0.053]**
P5→Subcortical 2y→Externalizing 2y: estimate = 0.024, 95% bootstrap CI = [0.0068, 0.046]

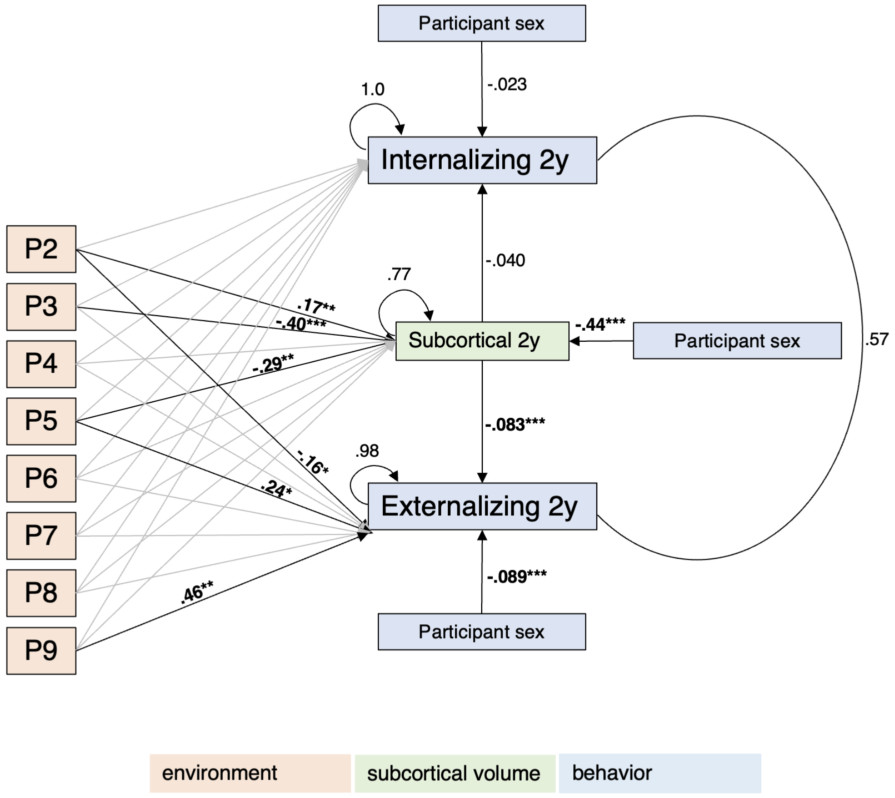


**Figure S3. Integrated approach linking multi-system environmental profiles, subcortical GM volume, and externalizing/internalizing psychopathology.** The integrated approach was operationalized using a path analysis from structural equation modeling and participant sex assigned at birth was treated as a covariate. The multi-system environmental profiles (denoted by P2, …., P9) are derived from family, school, neighborhood, and policy factors at baseline. The imaging measure corresponds to the subcortical gray matter (GM) volume at 2-year follow-up. Externalizing and internalizing psychopathology are derived from the Child Behavior Checklist (CBCL) at 2-year follow-up. Bold black arrows represent the significant relationships (direct and indirect) with their respective coefficient estimates. Gray arrows indicate the relationships between the multi-level environmental profiles and subcortical GM volume or externalizing/internalizing psychopathology that are not significant. Circle arrows capture the respective variances of subcortical GM volume, externalizing psychopathology, and internalizing psychopathology. Circular line denotes the correlation between externalizing and internalizing psychopathology. A bootstrapping procedure was performed to estimate the 95% confidence intervals (CIs) of the indirect effects by selecting samples of youth randomly to perform the path analysis over 10,000 times. *$P < 0.05$. **$P < 0.01$. ***$P < 0.001$.
